# A map of canine sequence variation relative to a Greenland wolf outgroup

**DOI:** 10.1101/2024.06.03.597150

**Authors:** Anthony K. Nguyen, Peter Z. Schall, Jeffrey M. Kidd

## Abstract

For over 15 years, canine genetics research relied on a reference assembly from a Boxer breed dog named Tasha (i.e., canFam3.1). Recent advances in long-read sequencing and genome assembly have led to the development of numerous high-quality assemblies from diverse canines. These assemblies represent notable improvements in completeness, contiguity, and the representation of gene promoters and gene models. Although genome graph and pan-genome approaches have promise, most genetic analyses in canines rely upon the mapping of Illumina sequencing reads to a single reference. The Dog10K consortium, and others, have generated deep catalogs of genetic variation through an alignment of Illumina sequencing reads to a reference genome obtained from a German Shepherd Dog named Mischka (i.e., canFam4, UU_Cfam_GSD_1.0). However, alignment to a breed-derived genome may introduce bias in genotype calling across samples. Since the use of an outgroup reference genome may remove this effect, we have reprocessed 1,929 samples analyzed by the Dog10K consortium using a Greenland wolf (mCanLor1.2) as the reference. We efficiently performed remapping and variant calling using a GPU-implementation of common analysis tools. The resulting call set removes the variability in genetic differences seen across samples and breed relationships revealed by principal component analysis are not affected by the choice of reference genome. Using this sequence data, we inferred the history of population sizes and found that village dog populations experienced a 9-13 fold reduction in historic effective population size relative to wolves.

## Introduction

For almost two decades the canine genome research community relied on a reference genome derived from a Boxer breed dog named Tasha (Lindblad-Toh et al. 2005). This genome was assembled using a Sanger capillary sequencing whole genome shotgun approach coupled with limited finishing of selected BAC clones. The Boxer-derived genome served as a reference for gene annotation, for the discovery of genetic variation across canines, and the development of canine SNP genotyping arrays (Vaysse et al. 2011). The assembly was further improved through targeted finishing and gene annotation efforts (Hoeppner et al. 2014). The resulting genome, known as canFam3.1 based on its description in the UCSC Genome Browser, became the standard reference for canine genomics (Nassar et al. 2023). With the advent of routine whole genome sequencing using Illumina short-read technology, an increasing number of canine genomes were sequenced and aligned to this Boxer-derived reference (Jagannathan et al. 2019; Plassais et al. 2019). This included ancient DNA data from canines, which involves sequencing of short, damaged DNA fragments, which has refined the understanding of dog origins (Bergstrom et al. 2020; Bergstrom et al. 2022; Botigue et al. 2017; Frantz et al. 2016; Ni Leathlobhair et al. 2018; Ramos-Madrigal et al. 2021; Skoglund et al. 2015).

Although bias due to the ascertainment of SNPs that are genotyped on arrays is well known (Clark et al. 2005; Lachance and Tishkoff 2013), it is not clear how reliance on a single reference may affect genetic variation studies based on sequencing. A systematic bias toward reporting the reference allele has been reported previously, particularly when analyzing short-read data associated with ancient DNA (Gunther and Nettelblad 2019; Günther and Schraiber 2024; Martiniano et al. 2020; Orlando et al. 2013; Swiel et al. 2024) or allele-specific RNA-sequencing analysis (Stevenson et al. 2013). In 2017 Gopalakrishnan and colleagues reported the assembly of the genome of a Swedish wolf using Illumina short-read sequencing data (Gopalakrishnan et al. 2017). Although the resulting assembly was highly fragmented, with a contig N_50_ of 94 kbp, it captured most of the unique sequence present in canines. Reanalysis of existing dog and wolf short-read data using this reference revealed patterns of variation highly similar to those found when using the Boxer-derived genome reference. However, the authors noted that dogs showed a larger estimated heterozygosity when aligned to the canFam3.1 assembly as opposed to the Swedish wolf assembly. The contribution of assembly quality to this metric is unclear.

Soon after the release of the fragmented Swedish wolf genome, the development of long-read sequencing technologies revolutionized genomics. This led to the release of multiple high-quality reference assemblies from canines, including several different breed dogs (Edwards et al. 2021; Field et al. 2020; Field et al. 2022; Halo et al. 2021; Jagannathan et al. 2021; Player et al. 2021; Schall et al. 2023; Sinding et al. 2021; Wang et al. 2021). These assemblies have high accuracy and continuity, revealing new features that were absent from the canFam3.1 assembly such as GC-rich segments, promoters, and the first exons of gene models. In 2023, the Dog10K consortium released a systematic analysis of canine genome variation based on whole genome sequence data from 1,929 dogs and wolves aligned to the UU_Cfam_GSD_1.0/canFam4 assembly derived from a German Shepherd Dog (Meadows et al. 2023). This included 57 wolves from across Eurasia, 281 village dogs sampled from 26 different countries, 1,579 dogs from 321 breeds, and 12 dogs with a mixed origin or that are not recognized by any international registering body. The Dog10K project represents the largest-scale genome analysis performed in dogs using a reference other than the Boxer-derived canFam3.1 assembly.

Among the new suite of long-read reference genomes is mCanLor1.2, which is derived from a Greenland wolf (Sinding et al. 2021). This wolf sample was chosen by the Darwin Tree of Life Project as a close outgroup to dogs and Eurasian wolves with the least coyote admixture. The Greenland wolf belongs to the polar wolf group in North America and can serve as an outgroup to dogs, which have an origin in Eurasia (Sinding et al. 2018). The assembly was constructed using Pacific Biosciences HiFi circular consensus reads, a sequencing technology that creates highly-accurate long reads yielding higher quality assemblies compared to previous methods (Wenger et al. 2019). The HiFi read data was further supplemented with Hi-C and 10X Genomics Read Cloud Sequence data to generate a chromosome-scale assembly following methods developed by the Vertebrate Genomes Project (Rhie et al. 2021) and The Darwin Tree of Life Project (Darwin Tree of Life Project 2022). The resulting chromosome-scale scaffolds were assigned to named chromosomes based on synteny with an existing Labrador retriever genome assembly (GCF_014441545.1).

In this study, we take advantage of recent informatics advances that use the highly parallel processing capabilities of graphics processing units (GPUs) to reanalyze whole genome sequencing data from 1,929 dogs and wolves sequenced by the Dog10K consortium using the mCanLor1.2 Greenland wolf genome as a reference. We first compare the structure of the mCanLor1.2 and UU_Cfam_GSD_1.0 assemblies, then describe patterns of SNP variation found when comparing to each assembly. Finally, we use the wolf-aligned data set to infer the history of population size changes in populations of village dogs and wolves.

## Results

### Global comparison of mCanLor1.2 and UU_Cfam_GSD_1.0 assemblies

Our previous analysis confirmed the high contiguity and completeness of the mCanLor1.2 assembly relative to other recently published canine assemblies and confirmed that mCanLor1.2 is an outgroup relative to the variation present in assemblies from breed dogs and a Basenji (Nguyen et al. 2024; Schall et al. 2023). Alignment of mCanLor1.2 and UU_Cfam_GSD_1.0 confirmed the large-scale continuity between the assemblies but revealed that the mCanLor1.2 chromosomal orientation is essentially random, with 19/39 analyzed chromosomes represented in the opposite orientation relative to UU_Cfam_GSD_1.0 (Figure 1). Consistent with the inverted orientation of the mCanLor1.2 X chromosome, analysis of read-depth profiles from male and female samples identified the mCanLor1.2 pseudo autosomal region (PAR) as chrX:117984001-124665963 (Figure S1).

**Fig. 1.**
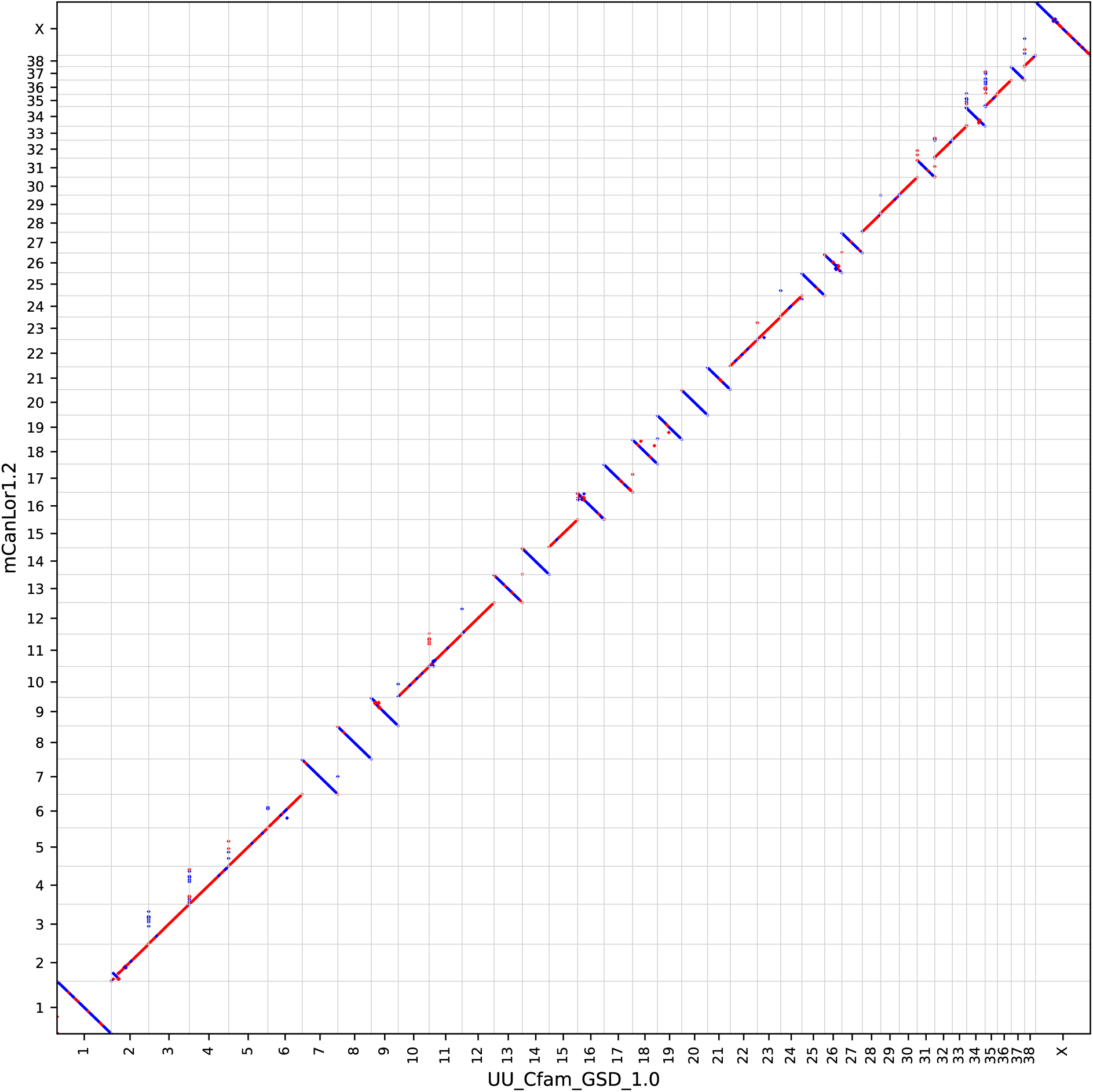
Chromosomal continuity between UU_Cfam_GSD_1.0 and mCanLor1.2. Corresponding segments between the UU_Cfam_GSD_1.0 (x axis) and mCanLor1.2 (y axis) assemblies were identified using minimap2. The position of aligned segments at least 5,000 bp in length are plotted. Alignments in the forward orientation are shown in red, alignments in the reverse orientation are shown in blue.

The mCanLor1.2 assembly was derived from a male sample and contains a partial assembly of the wolf Y chromosome. The sequence labeled as chrY (HG994382.1) consists of 6.5 Mbp of ampliconic sequence in a palindromic configuration (Figure S2). Three additional unlocalized sequences associated with the Y chromosome (CAJNRB020000005.1, CAJNRB020000035.1, and CAJNRB020000036.1) show homology to KP081776.1, a partial dog Y chromosome assembly (Li et al. 2013). The unique portions of KP081776.1, which have been previously used as a reference to study Y chromosome sequence variation (Oetjens et al. 2018; Smeds et al. 2019), largely correspond to portions of CAJNRB020000035.1 and CAJNRB020000036.1. Read-depth analysis of Illumina sequencing reads from the same sample used to generate mCanLor1.2 indicates that CAJNRB020000005.1, CAJNRB020000035.1, and CAJNRB020000036.1 are composed of mixtures of amplified and unique sequence, and that there are additional copies of Y-chromosome palindromes that remain unassembled (Figure S3).

We have previously compared the duplication content of existing canine genome assemblies, finding that the mCanLor1.2 assembly shows the highest concordance between read-depth and assembly self-alignment approaches (Nguyen et al. 2024). To further assess the mCanLor1.2 assembly, we aligned it to UU_Cfam_GSD_1.0 and identified 35,192 segments spanning 18,102,207 bp that are missing in UU_Cfam_GSD_1.0 but present in mCanLor1.2. This includes 9,003 insertions (1,836,394 bp) associated with SINEC sequences and 1,462 (613,486 bp) associated with LINE-1 sequences. Of the 35,192 segments that are missing in UU_Cfam_GSD_1.0, we identified 418 that overlap with an exon annotated in ENSEMBL rapid release mCanLor1.2 annotation (Table S1).

### Identification of Canine Sequence Variation Relative to mCanLor1.2

Using the GPU-accelerated pipeline implemented in the NVIDIA Clara Parabricks software (Franke and Crowgey 2020), we aligned reads from 1,929 dogs and wolves to the mCanLor1.2 assembly and identified variants using the same procedures previously used by the Dog10K platform. We identified a total of 36,908,138 SNVs and 13,640,837 indels. This represents an ∼6.7% increase over the 34,566,550 SNVs identified based on alignment to UU_Cfam_GSD_1.0. We calculated the distance between each sample and the reference genome as half the number of heterozygous genotypes plus the number of homozygous alternative genotypes.

When using the UU_Cfam_GSD_1.0 reference, which was derived from a German Shepherd Dog, the total distance from the UU_Cfam_GSD_1.0 reference varies widely across sample groups (Figure 2). Breed dogs range from 1,470,039 to 2,908,571 differences (median 2,381,059), village dogs range from 1,689,172 to 2,899,436 differences (median 2,535,491) and wolves range from 3,471,282 to 3,582,520.5 differences (median 3,546,063). The range of divergence among breed and village dogs is drastically reduced when using the mCanLor1.2 Greenland wolf. In this call set, breed dogs range from 3,720,950.5 to 3,803,669 differences (median 3785952.5), village dogs range from 3,720,129 to 3,817,528 (median 3,784,434.5) and wolves range from 3,563,551 to 3,810,647 (median 3,706,174.5). Although the total number of variants differs greatly based on the references employed, the number of heterozygous SNV genotypes identified in each sample shows a strong correlation between the call sets (Spearmen r = 0.9999, Figure S4). Principal component analysis recapitulates the sample relationships found when using UU_Cfam_GSD_1.0 as the reference. For example, when analyzing a single sample from each of 314 breeds, both SNV sets separate German Shepherds and related breeds from other samples along Principal Component 3 (Figure 3). Thus, the observed sample projections do not reflect a bias induced by alignment to a reference derived from a German Shepherd Dog.

**Fig 2.**
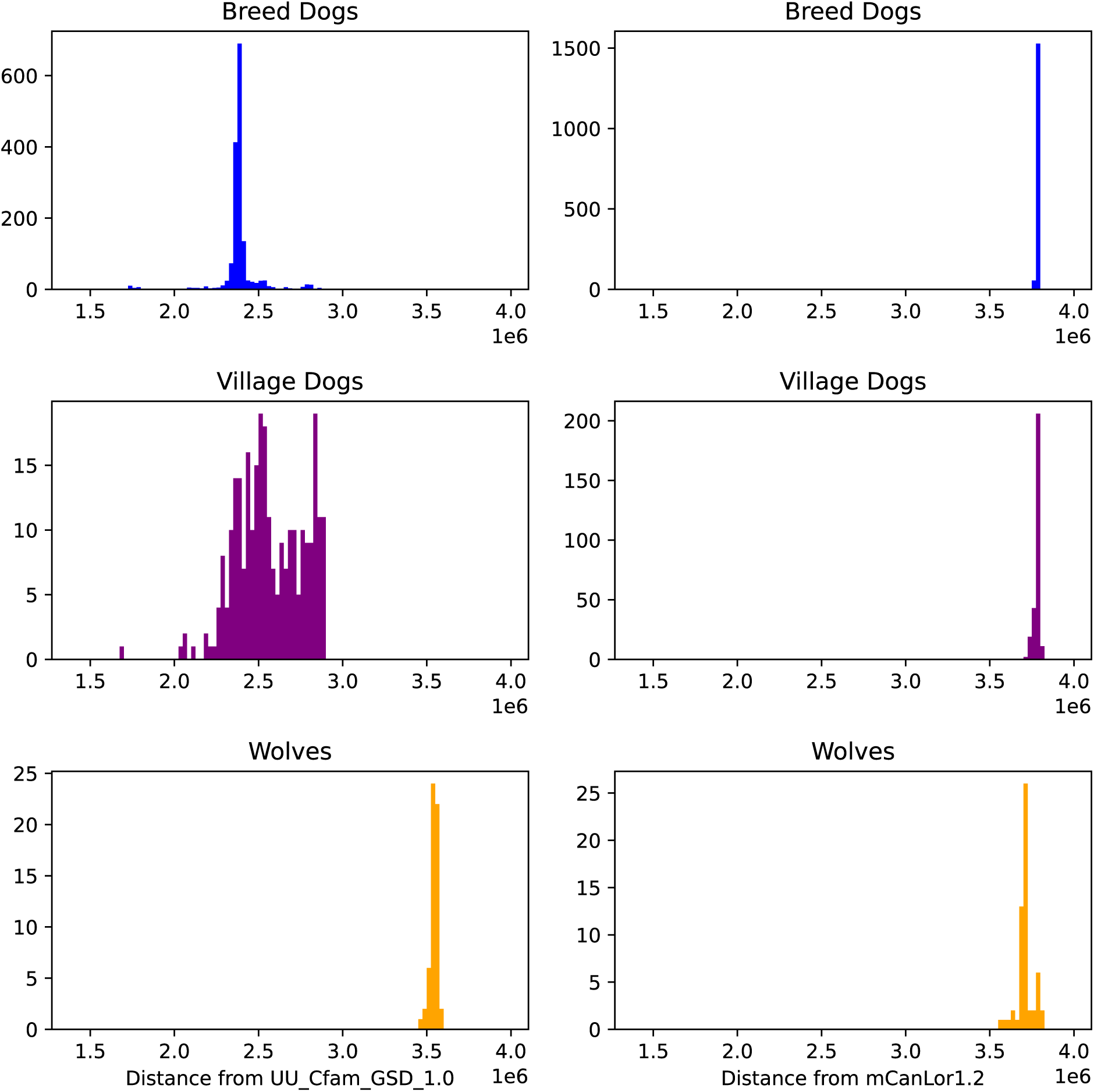
Distance between sequenced samples and the UU_Cfam_GSD1.0 and mCanLor1.2 references. The distance between each sample and the reference genome was calculated as half the number of heterozygous genotypes plus the number of homozygous alternative genotypes. Histograms of the distance between each sample and each reference are shown with breed dogs, village dogs, and wolves plotted separately.

**Fig 3.**
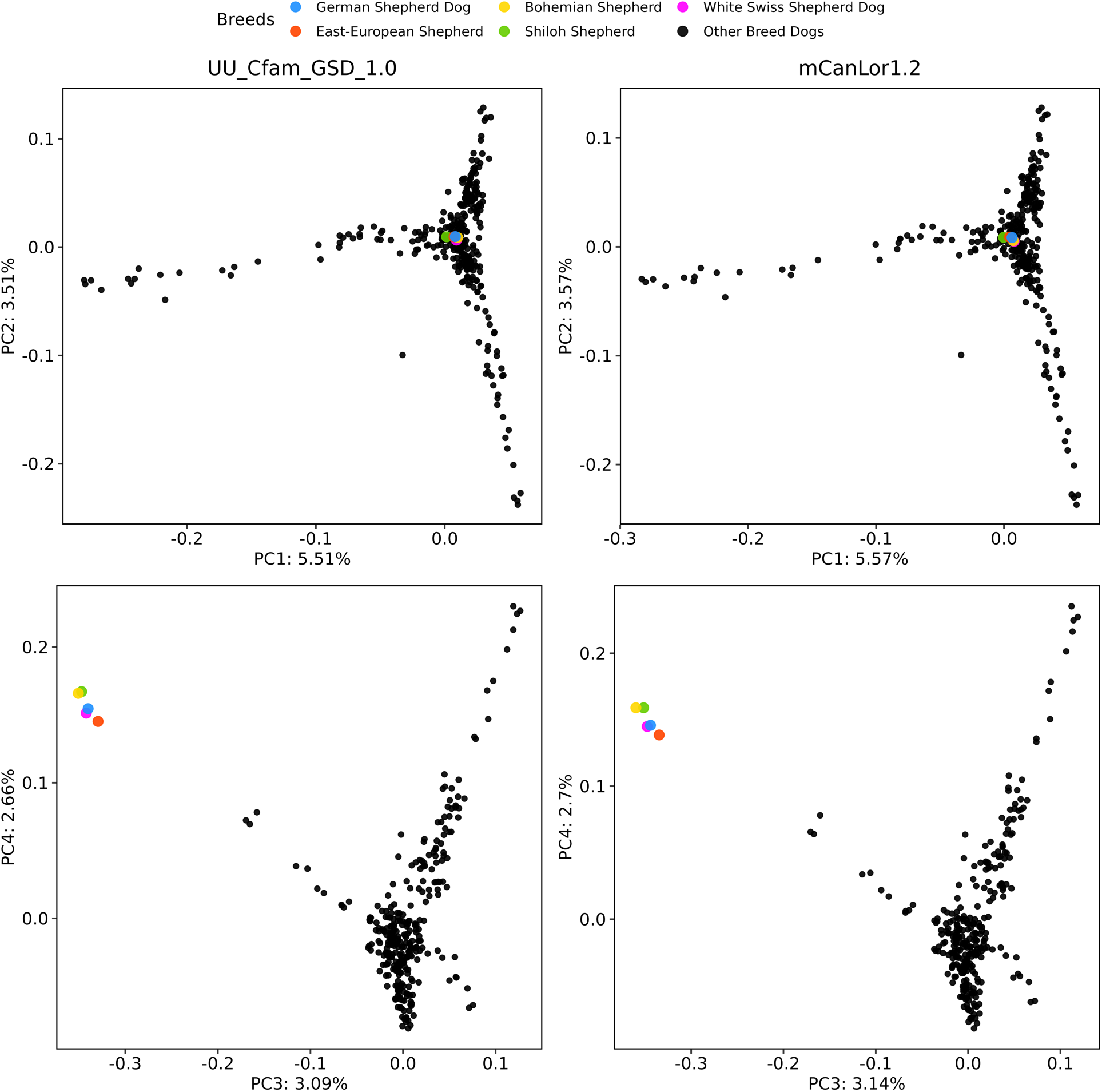
Principal Component Analysis of Breed Dogs is Similar Using Both References. Principal component analysis was performed using one sample from each of 314 breeds based on the variants derived from alignment to the UU_Cfam_GSD_1.0 (left) and mCanLor1.2 (right) assemblies. In both cases, projections of samples onto the 1^st^ vs 2^nd^ and 3^rd^ vs 4^th^ components are shown. Breeds related to German Shepherd Dogs are colored as indicated and separate from the other breeds along principal component 3.

### Estimates of Village Dog and Wolf Demographic Histories

Using the alignments to the mCanLor1.2 assembly, we estimated the trajectory of effective population size (N_E_) through time for four village dog and four wolf populations (Figure 4). The sampled village dogs and wolf populations do not share a similar population size trajectory until approximately 100,000 years ago, an observation consistent with findings from ancient DNA that indicate that a portion of dog genetic ancestry remained partially differentiated from sampled ancient wolves since before 100 kya (Bergstrom et al. 2022). From 20 – 40 thousand years ago village dog populations experienced a 9-13 fold reduction in effective population size. To explore the effects of the reference genome on these estimates, we repeated this analysis with one wolf and one village dog population using variants discovered using alignment to the UU_Cfam_GSD_1.0 German Shepherd Dog assembly (Fig S5). The profile obtained for village dogs from Congo is highly similar throughout the entire analyzed period while the inferred population size history for wolves from Greece is concordant for times older than 10,000 years and shifted slightly for more recent time periods.

**Fig 4.**
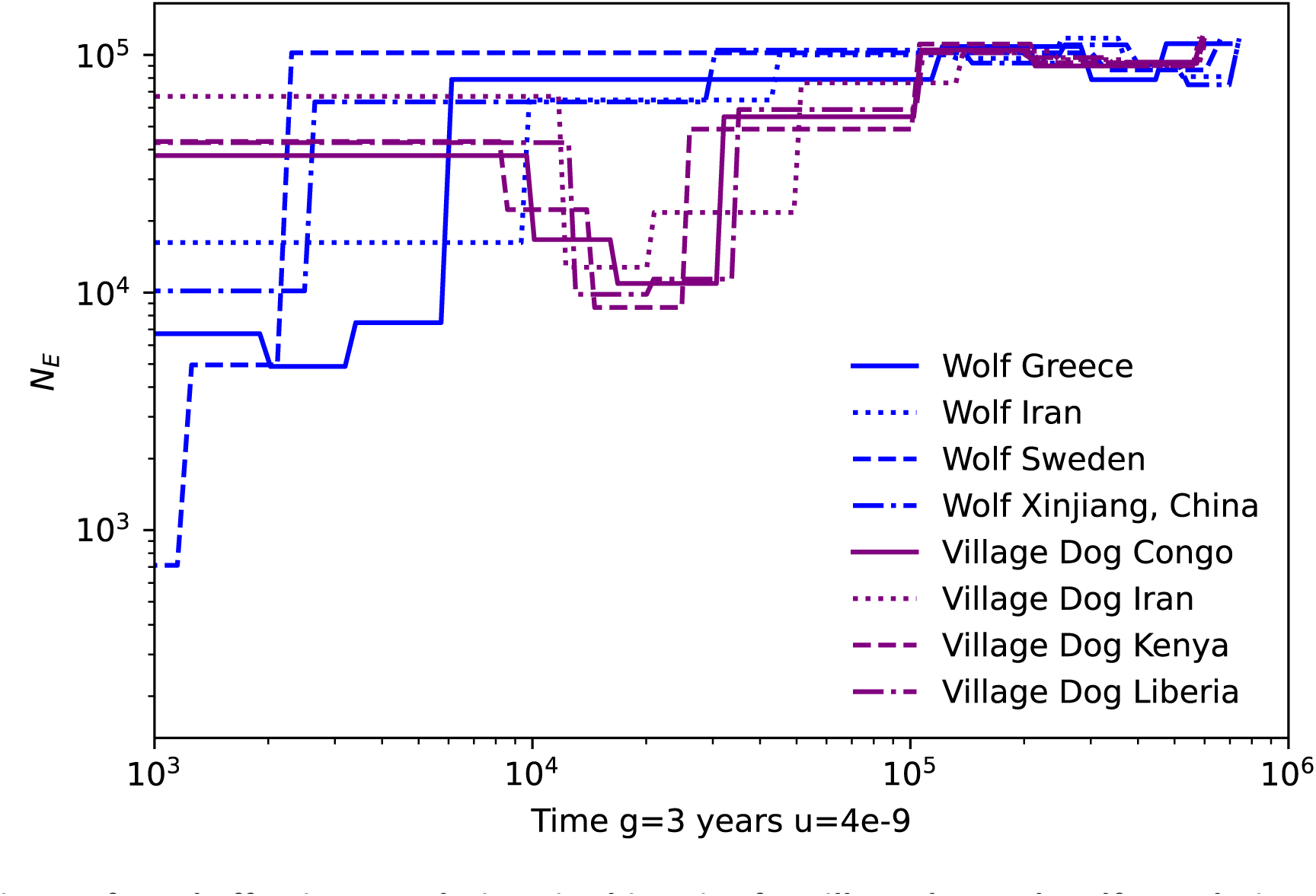
Inferred effective population size histories for village dog and wolf populations. The population history of four wolf populations (blue lines: Greece (n=12), Iran (n=6), Sweden (n=6), and Xinjiang, China (n=6)) and four village dog populations (purple lines: Congo (n=16), Iran (n=24), Kenya (n=18), and Liberia (n=17)) were inferred using SMC++. Values were plotted assuming a generation time of 3 years and a mutation rate of 4 * 10^-9^ per site per generation.

## Discussion

The advent of efficient technologies for genome assembly have enabled graph-based pan genome analyses (Eizenga et al. 2020; Gong et al. 2023). Despite the promise of these approaches, alignment of short-read genome sequence data to a single reference genome remains the dominant paradigm for studies of genetic variation. This may introduce a bias in comparisons among samples since sequencing reads that are more similar to the genome reference sequence are more likely to align. Although careful consideration of alignment tools and parameters can ameliorate this effect (Oliva et al. 2021), the potential effects of the reference on downstream analyses are largely unexplored. Due to the complex history of inbreeding that underlies the formation of modern dog breeds (Parker et al. 2017), the reliance on a single reference genome derived from a particular breed may bias the analysis of genetic differences within and among canines.

This potential bias can be overcome using an appropriate outgroup species as the reference. Recent advances in long-read genome sequencing and assembly approaches have led to the publication of multiple new canine reference genomes. However, reanalyzing existing genome-wide data collections requires an onerous and computationally expensive process of remapping and variant calling. In this study we combined new, GPU-based implementations of standard alignment and variant identification pipelines to efficiently reanalyze whole genome sequence data from 1,929 dogs and wolves using a high-quality Greenland wolf genome as the reference.

A direct speed comparison between analysis pipelines is challenging since some implementations may be better tailored to the architecture of a particular computer cluster or more sensitive to file input-output load caused by other users. To provide an estimate of the relative improvement of the Parabricks GPU pipeline we reprocessed Illumina sequence data from a Cocker Spaniel breed dog using both a standard pipeline and the Parabricks pipeline. In this comparison, the Parabricks pipeline had an 8.8-fold improvement in run time (53 minutes vs 470 minutes, Fig S6).

As expected, use of the wolf genome normalizes the apparent divergence between dog samples and the reference. General patterns of genome diversity are unchanged when either the Greenland wolf or a German Shepherd Dog reference are employed, including PCA analysis that shows a separation of German Shepherd-related breeds from others. Thus, the choice of reference genome appears to have little effect on variation discovered using moderate coverage genomes from modern samples. Given the high degree of allele sharing reported among German Shepherd-related breeds and other breed clades (Meadows et al. 2023; Parker et al. 2017), the Greenland wolf-aligned data set may be a useful resource for studies of the origins and evolution of modern dogs, particularly in the analysis of low-coverage ancient DNA samples.

Methods based on the sequentially Markovian coalescent are powerful approaches for inferring past population sizes based on genome-wide sequencing data (Mather et al. 2020). Taking advantage of alignment to the highly contiguous mCanLor1.2 assembly, we used SMC++ to infer the history of village dog and wolf populations. Unlike other approaches, SMC++ uses multiple individuals and does not require variation data to be phased, resulting in more accurate estimates (Terhorst et al. 2017). Application of SMC++ separately to wolves (6-12 individuals per population) and village dogs (16-24 individuals per population) indicates that the ancestors of village dogs experienced a 9-13 fold reduction in effective population size 20-40 thousand years ago. This reduction likely reflects domestication from an unsampled wolf population (Bergstrom et al. 2020) and is similar, though slightly smaller, than the ∼16-fold reduction in effective population size previously estimated using a Dingo and a Basenji using the pairwise sequential Markovian coalescent (PSMC) inference method (Freedman et al. 2014; Li and Durbin 2011).

## Material and Methods

### Genome Assembly Comparisons

The mCanLor1.2 (GCA_905319855.2) Greenland wolf assembly (Sinding et al. 2021) was downloaded from NCBI. The UU_Cfam_GSD_1.0 German Shepherd Dog assembly (Wang et al. 2021) (‘Mischka’, GCA_011100685.1, canFam4) was downloaded from the UCSC Genome Browser (RRID:SCR_005780). To assess continuity, the UU_Cfam_GSD_1.0 genome was aligned to mCanLor1.2 using Minimap2 version 2.26 (Li 2018, 2021) with option -x asm10. The resulting alignments were parsed to retain aligned segments with a query and target sequence length of at least 5,000 bp.

The approximate location of the Pseudoautosomal Region (PAR) on the X chromosome was determined by examination of read depth profiles from 3 male (VILLCR000002, CLUPGR000011, and BOXR000005) and 1 female (CLUPSE000004) samples using mosdepth version 0.3.2 (RRID:SCR_018929) (Pedersen and Quinlan 2018) with a window size of 1 kbp.

To assess representation of the Y chromosome, four mCanLor1.2 segments were compared: chrY (HG994382.1) as well as three unlocalized sequences associated with the Y chromosome: CAJNRB020000005.1, CAJNRB020000035.1, and CAJNRB020000036.1. Self-similarity dotplots were generated used Gepard version 2.1 (Krumsiek et al. 2007) with a word size of 100 bp and ampliconic arms defined using iterative alignment with Minimap2 version 2.26. The mCanLor1.2 Y-chromosome sequences were compared with KP081776.1, an assembly of a portion of the dog Y chromosome generated by pooled sequencing of BAC clones from a Doberman (Li et al. 2013) that has been previously used as a reference to study Y chromosome sequence variation among canids using short-read sequencing (Oetjens et al. 2018; Smeds et al. 2019). The positions of the Y-chromosome SNVs used in (Oetjens et al. 2018) were converted to mCanLor1.2 coordinates by aligning 101 bp long fragments centered on each variant using Minimap2 version 2.26. Read-depth profiles based on the Illumina data generated as part of the mCanLor1.2 assembly were calculated using fastCN as previously described (Nguyen et al. 2024; Pendleton et al. 2018).

Insertions in mCanLor1.2 relative to UU_Cfam_GSD_1.0 were defined using SVIM-asm version 1.0.3 (Heller and Vingron 2021) based on Minimap2 alignments. Analysis was limited to the 38 assembled autosomes and the X chromosome. Putative SINE insertions were identified as insertions 150-250 bp in size that contained at least 90% SINE sequence. Potential LINE-1 insertions were not filtered by length but were required to have at least 90% of the insertion sequence annotated as LINE. Intersecting exons were identified using the ENSEMBL rapid release of the mCanLor1.2 GTF file (https://ftp.ensembl.org/pub/rapid-release/species/Canis_lupus/GCA_905319855.2/ensembl/geneset/2022_01/Canis_lupus-GCA_905319855.2-2022_01-genes.gtf.gz)

### Sequence Alignment and Variant Identification Relative to mCanLor1.2

We identified single nucleotide (SNV) and short insertion-deletion (indel) variants based on alignment of Illumina data to the mCanLor1.2 assembly. Our procedure began with the aligned CRAM files from 1,929 samples that passed quality control filters described by the Dog10K consortium (Meadows et al. 2023). We note that the base qualities in the CRAM files were already recalibrated based on alignment to the UU_Cfam_GSD_1.0 assembly. First, reads from the CRAM files were converted to fastq format using the collate and bam2fq commands from SAMtools version 1.13 (RRID:SCR_002105). The resulting reads were then processed using the fq2bam and haplotypecaller commands from the GPU-accelerated NVIDIA Clara Parabricks tool kit version 4.0.1-1 (Franke and Crowgey 2020). This resulted in a genomic variant call format (GVCF) file for each sample.

Sequence variants were then identified from the 1,929 GVCF files following the same procedures previously used by the Dog10K Consortium (Meadows et al. 2023). First, candidate variants were identified using the *GenomicsDBImport* and *GenotypeGVCFs* commands in GATK version 4.2.0.0 (RRID:SCR_001876). SNV loci were separated from loci containing indels using the GATK *SelectVariants* command. High-quality SNVs were identified using the variant quality score recalibration (VQSR) procedure using 680,458 SNPs from the Illumina Canine HD and Axiom Canine HD SNP arrays as a truth set. Positions were converted from UU_Cfam_GSD_1.0 to mCanLor1.2 coordinates using the GATK *LiftoverVcf* utility with a genome chain file generated constructed from BLAT alignments (Hinrichs et al. 2006; Kent 2002). Due to differences in read depth profiles, variants on the autosomes and X-par region were analyzed separately from variants on the non-PAR portion of the X chromosome. In each case, cut offs were used such that 99.0% of variants in the truth set were retained, resulting in a total of 36,908,138 PASS SNVs for downstream analysis. Due to the absence of suitable training data, candidate indels were identified using the GATK *VariantFiltration* utility with the following criteria: QD < 2.0, FS > 200.0, *ReadPosRankSum* < -2.0, SOR > 10.0.

### SNV analysis

The distance between each sample and the reference genome was calculated as the number of homozygous alternative SNV genotypes plus half the number of heterozygous genotypes. Principal Component Analysis (PCA) was performed using *smartpca* from the EIGENSOFT package (version 8.0.0 RRID:SCR_004965) (Patterson et al. 2006). First, samples were filtered to retain only one sample from each of 314 breeds, keeping the sample with the highest coverage in the original Dog10K data set. Breeds with large amounts of recent wolf ancestry (Czechoslovakian Wolfdog, Saarloos Wolfdog, and Tamaskan) were omitted. Next, biallelic autosomal SNVs with a minor allele frequency of at least 0.05 and a genotyping missing rate less than 0.10 were selected using PLINK version 1.9 (RRID:SCR_000084) (Chang et al. 2015). The resulting variants were pruned for linkage disequilibrium using PLINK with the command --indep-pairwise 50 10 0.1.

Trajectories of effective population size (N_E_) through time were constructed using SMC++ (Terhorst et al. 2017) version v1.15.4.dev18+gca077da.d20210316. Analysis was performed on eight groups including 12 wolves from Greece, 6 wolves from Iran, 6 wolves from Sweden, 6 wolves form Xinjiang, China, 16 village dogs from Congo, 24 village dogs from Iran, 18 village dogs from Kenya, and 17 village dogs from Liberia. Variation within regions of the mCanLor12 assembly identified as segmental duplications was removed using the *–mask* option in SMC++ (Nguyen et al. 2024). In each case, analysis was performed for each autosome and was repeated using five different distinguished individuals, based on the lexicographical order of sample identifiers. Results were converted from coalescent units using a generation time of 3 years and a mutation rate of 4 * 10^-9^ per site per generation, which is compatible with estimates from ancient DNA (Skoglund et al. 2015) and wolf pedigrees (Koch et al. 2019). For comparison, effective population size trajectories were also inferred for wolves from Greece and village dogs from Congo using variant calls based on alignment to the UU_Cfam_GSD_1.0 assembly produced by the Dog10K consortium.

### Run time comparison

To estimate the relative run time improvement of the NVIDIA Clara Parabricks tool kit we performed a reanalysis of ACKR000001, a Cocker Spaniel breed dog. The original reads (SRR12330404) were aligned to the UU_Cfam_GSD_1.0 assembly with the standard pipeline used by the Dog10K consortium (Meadows et al. 2023)(https://github.com/jmkidd/dogmap). This pipeline uses bwa-mem2 (Vasimuddin et al. 2019) (version 2.2.1) and GATK (McKenna et al. 2010) (version 4.2.0.0). Analysis was also performed using the Parabricks tool kit. Analysis was repeated three times using each approach. The standard CPU pipeline ran on a single compute node with an attached solid-state drive (SSD) and used 24 3.0 GHz Intel Xeon Gold 6154 CPUs. The Parabricks pipeline ran on a single compute node and used two NVIDIA Tesla V100 GPUs and 24 2.4 GHz Intel Xeon Gold 6148 CPUs.

## Supplementary Information

The online version contains supplementary material including Supplementary Table 1 and Supplementary Figures 1-6.

## Acknowledgements

We thank all members of the Dog10K consortium for contributing their samples and data to the project and the University of Michigan Advanced Research Computing Group for assistance with efficient GPU utilization. This research was supported in part through computational resources and services provided by Advanced Research Computing at the University of Michigan, Ann Arbor.

## Data Availability

SNV and indel variant data is available from the Zenodo archive under DOI 10.5281/zenodo.11248171.

## Competing Interests

The authors declare that no competing interests exist.

## Supplementary Figures

**Fig. S1.**
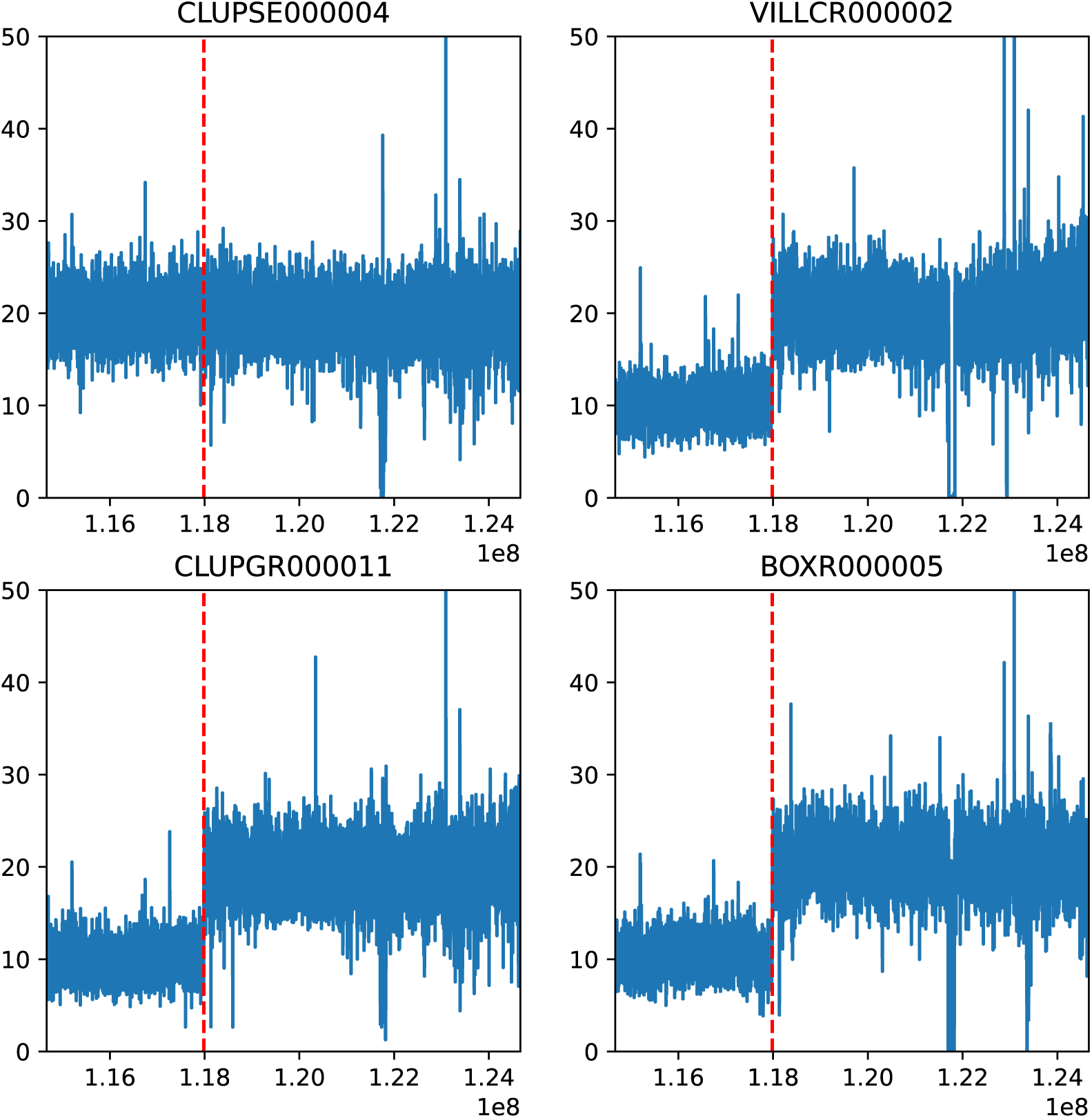
Identification of the mCanLor1.2 pseudo autosomal region. Read depth profiles are shown for 1 female (CLUPSE000004) and 3 male (VILLCR000002, CLUPGR000011, and BOXR000005) samples along the end of the X chromosome. The dashed red line represents the inferred PAR boundary, which extends from chrX:117984001-124665963.

**Fig S2.**
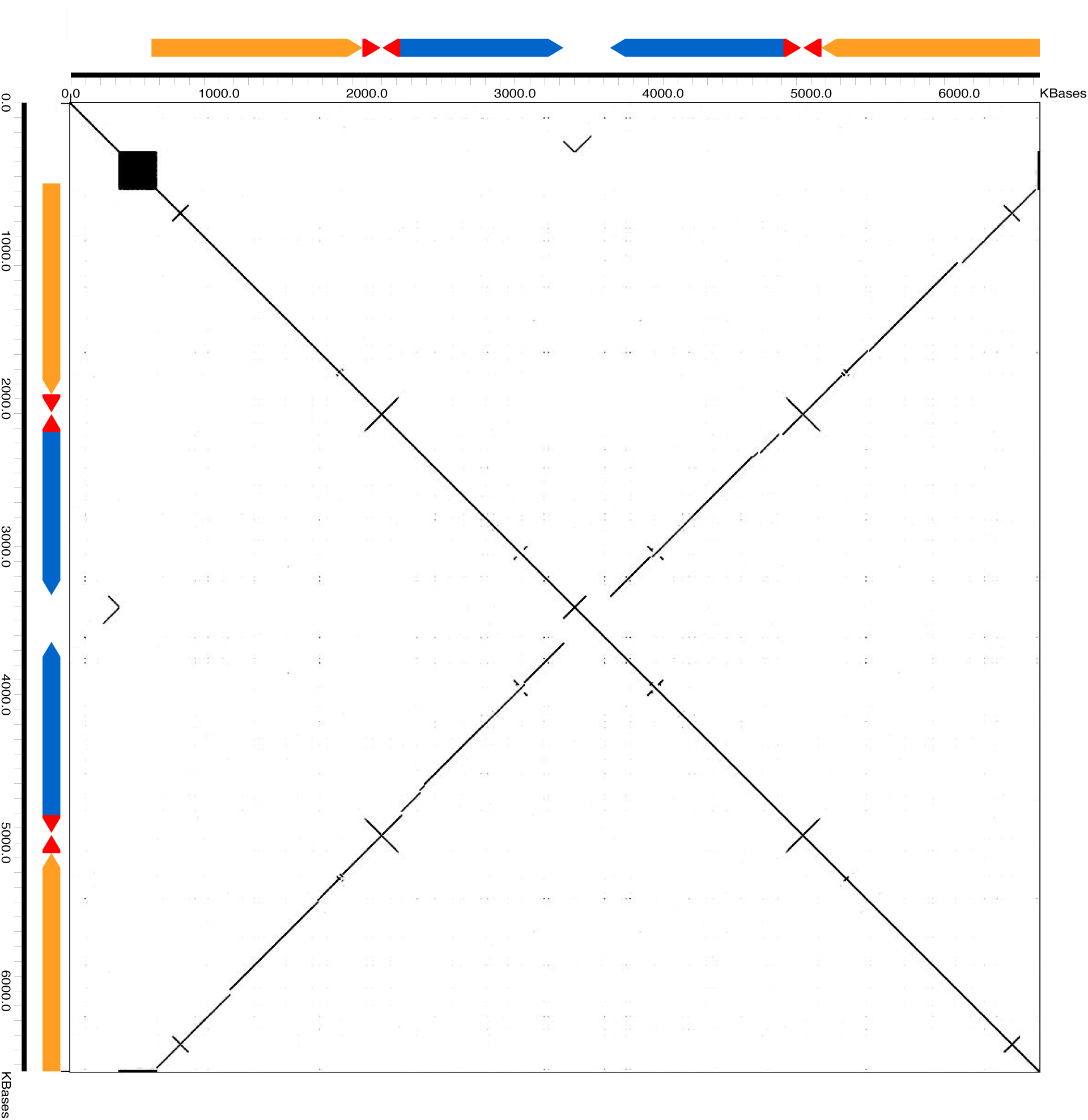
Ampliconic structure of chrY (HG994382.1) A self-similarity dotplot of the mCanLor1.2 Y chromosome sequence drawn with a window size of 100 bp is shown. The colored arrows correspond to ampliconic sequences identified in the assembly. The entire sequence represents a complex palindromic structure.

**Fig S3.**
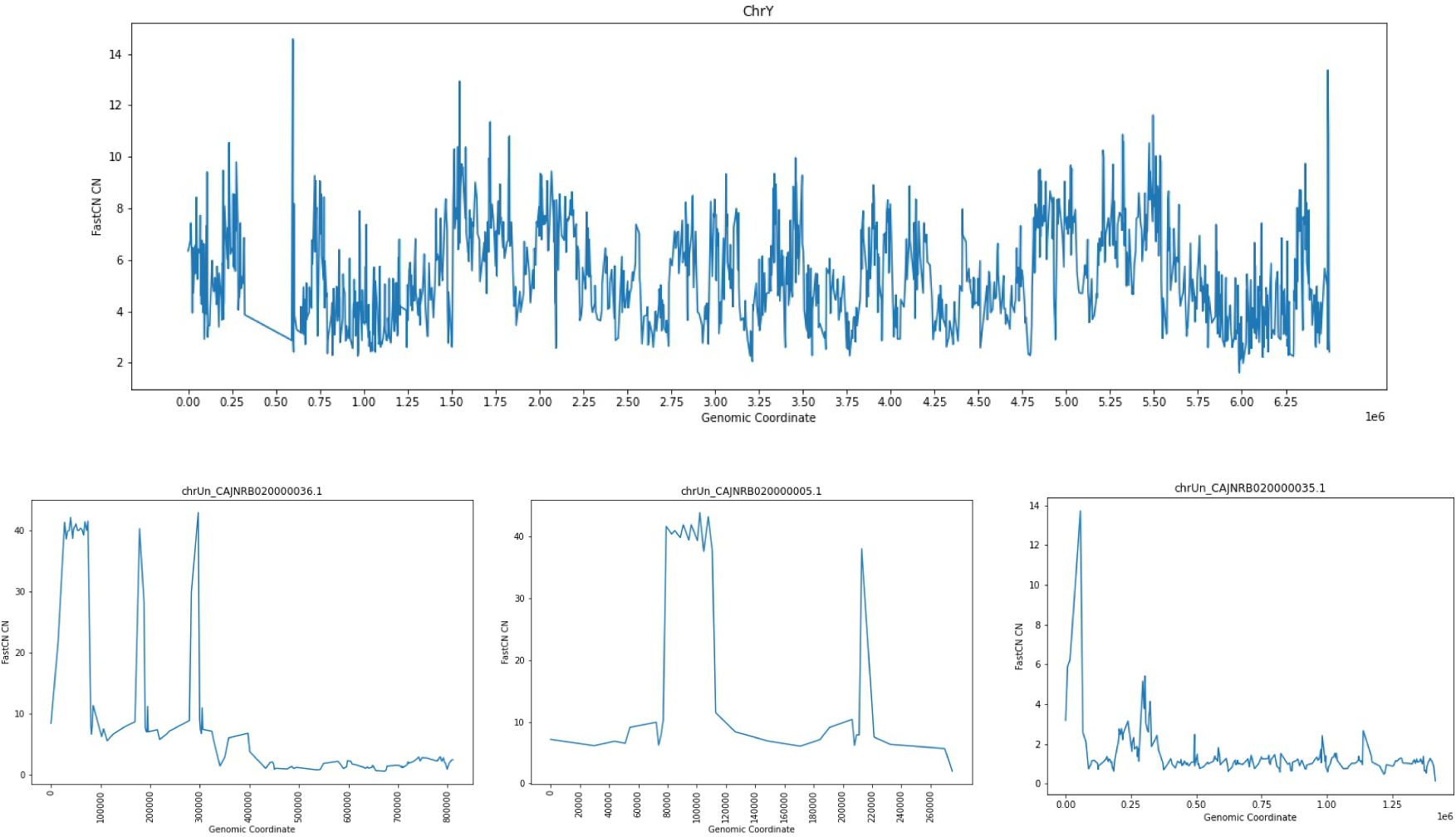
Copy-number profile across Y-chromosome sequences. Copy-number was inferred using fastCN applied to Illumina genome sequence data generated from the sample used to create the mCanLor1.2 assembly. Results are shown for the mCanLor1.2 Y-chromosome sequence (top) as well as three additional unplaced assembled contigs that are associated with the Y chromosome.

**Fig S4.**
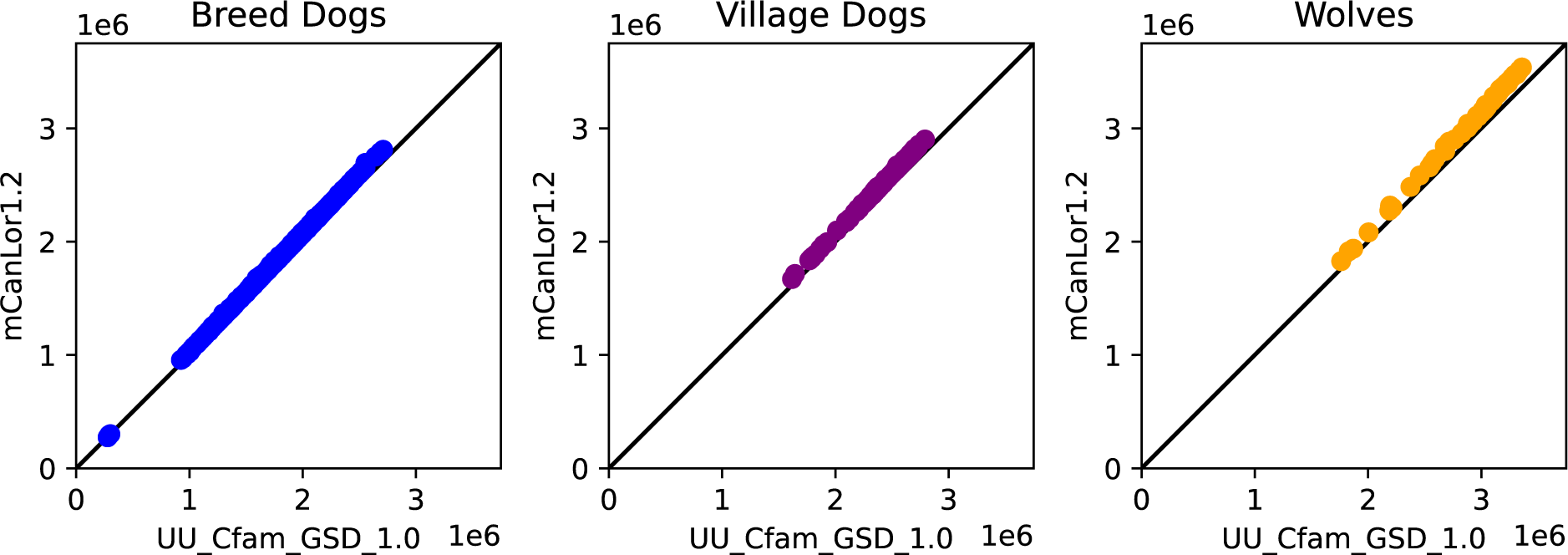
Number of heterozygous sites identified based on alignment to UU_Cfam_GSD_1.0 and mCanLor1.2. A scatter plot of the number of heterozygous sites identified from alignment to each reference genome is shown for breed dogs, village dogs, and wolves.

**Fig S5.**
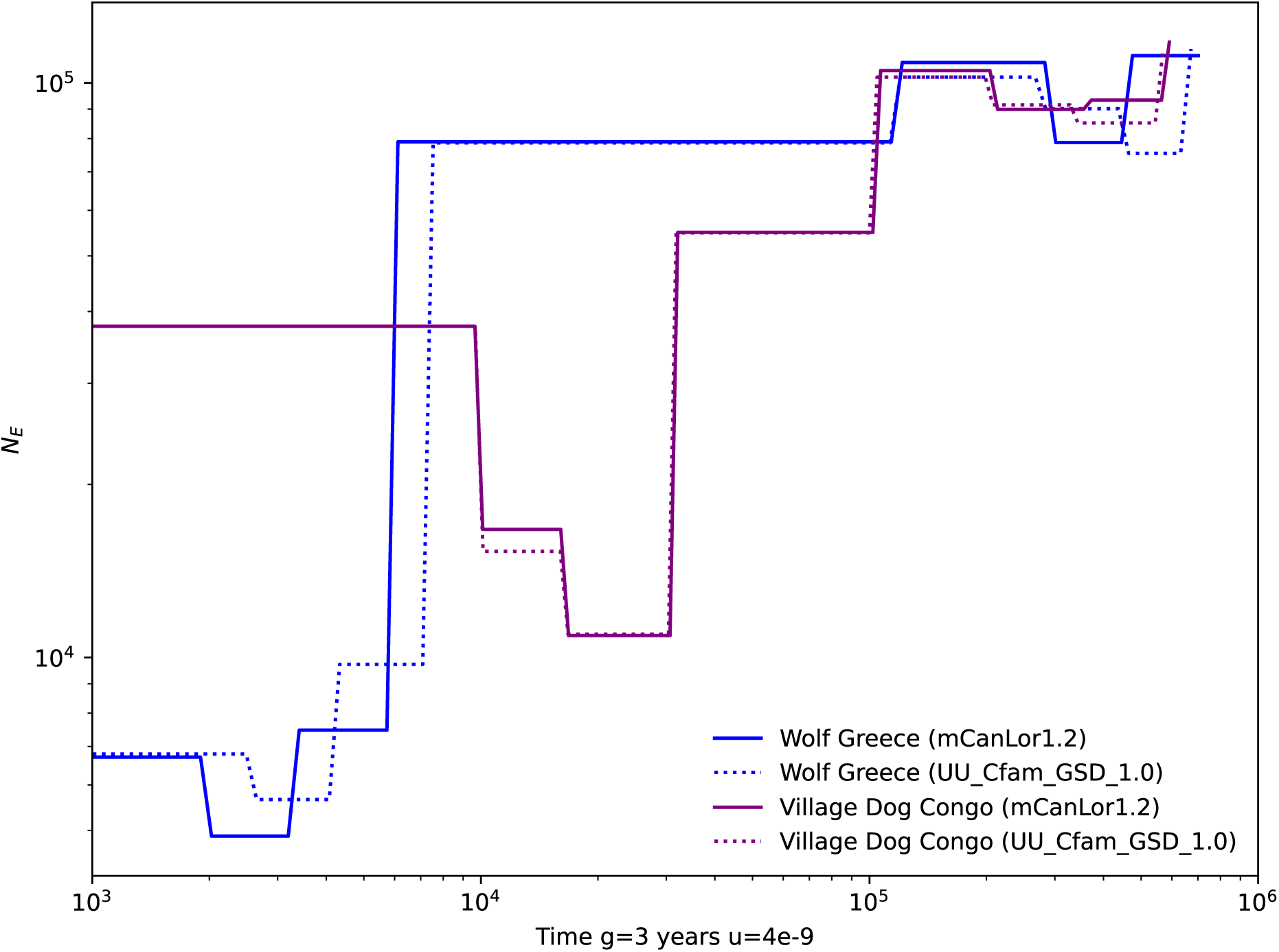
Comparison of population size histories inferred using different reference assemblies. The population history for wolves from Greece (blue lines) and village dogs from Congo (purple lines) was estimated using sequence alignment to the mCanLor1.2 (solid lines) and UU_Cfam_GSD_1.0 (dashed lines) assemblies. Inference was performed using SMC++.

**Fig S6.**
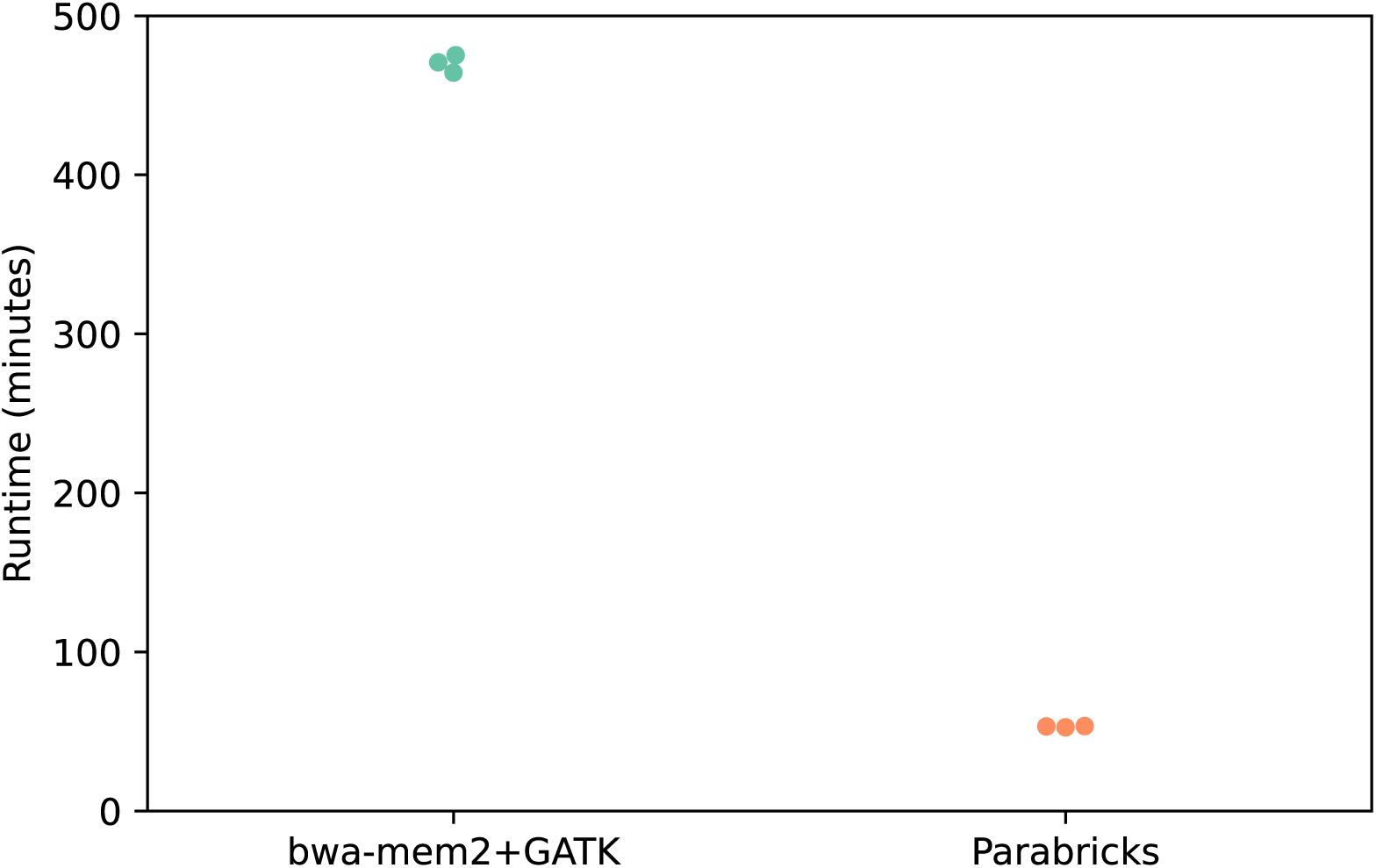
Runtime comparison of standard and GPU analysis pipelines. Illumina fastq reads from sample ACKR000001 (SRR12330404) were processed against the UU_Cfam_GSD_1.0 assembly using a standard pipeline (left) and the GPU-enabled Parabricks pipeline (right). Total runtime (wall time) results are plotted from three analysis runs. Analysis includes alignment to the reference genome, sorting and duplication marking, base recalibration, and generation of GVCF-formatted variant files. The standard CPU pipeline ran on a single compute node with an attached solid-state drive (SSD) and used 24 3.0 GHz Intel Xeon Gold 6154 CPUs. The Parabricks pipeline ran on a single compute node and used two NVIDIA Tesla V100 GPUs and 24 2.4 GHz Intel Xeon Gold 6148 CPUs.

**Supplementary Table 1 Sequence deleted in UU_Cfam_GSD_1.0 that overlaps with annotated exons** (see excel file)

## Notes

https://zenodo.org/doi/10.5281/zenodo.11248170

### Competing Interest Statement

The authors have declared no competing interest.

### Summary of Updates

Additional comparisons between variant sets and analysis of speed increase using GPUs.

