## Supplemental Figures for "A map of canine sequence variation relative to a Greenland wolf outgroup"

### Supplementary Figures

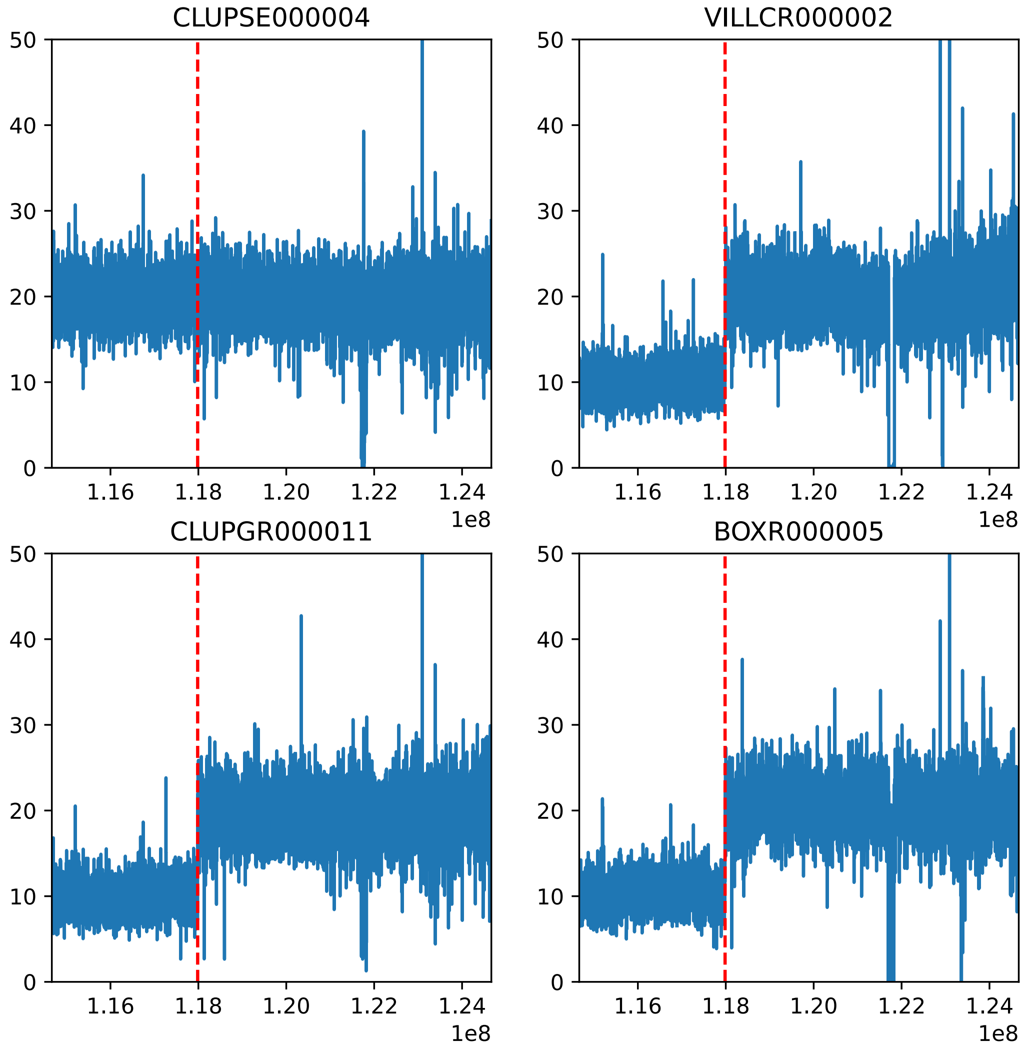


**Fig. S1. Identification of the mCanLor1.2 pseudo autosomal region**
Read depth profiles are shown for 1 female (CLUPSE000004) and 3 male (VILLCR000002, CLUPGR000011, and BOXR000005) samples along the end of the X chromosome. The dashed red line represents the inferred PAR boundary, which extends from chrX:117984001-124665963.

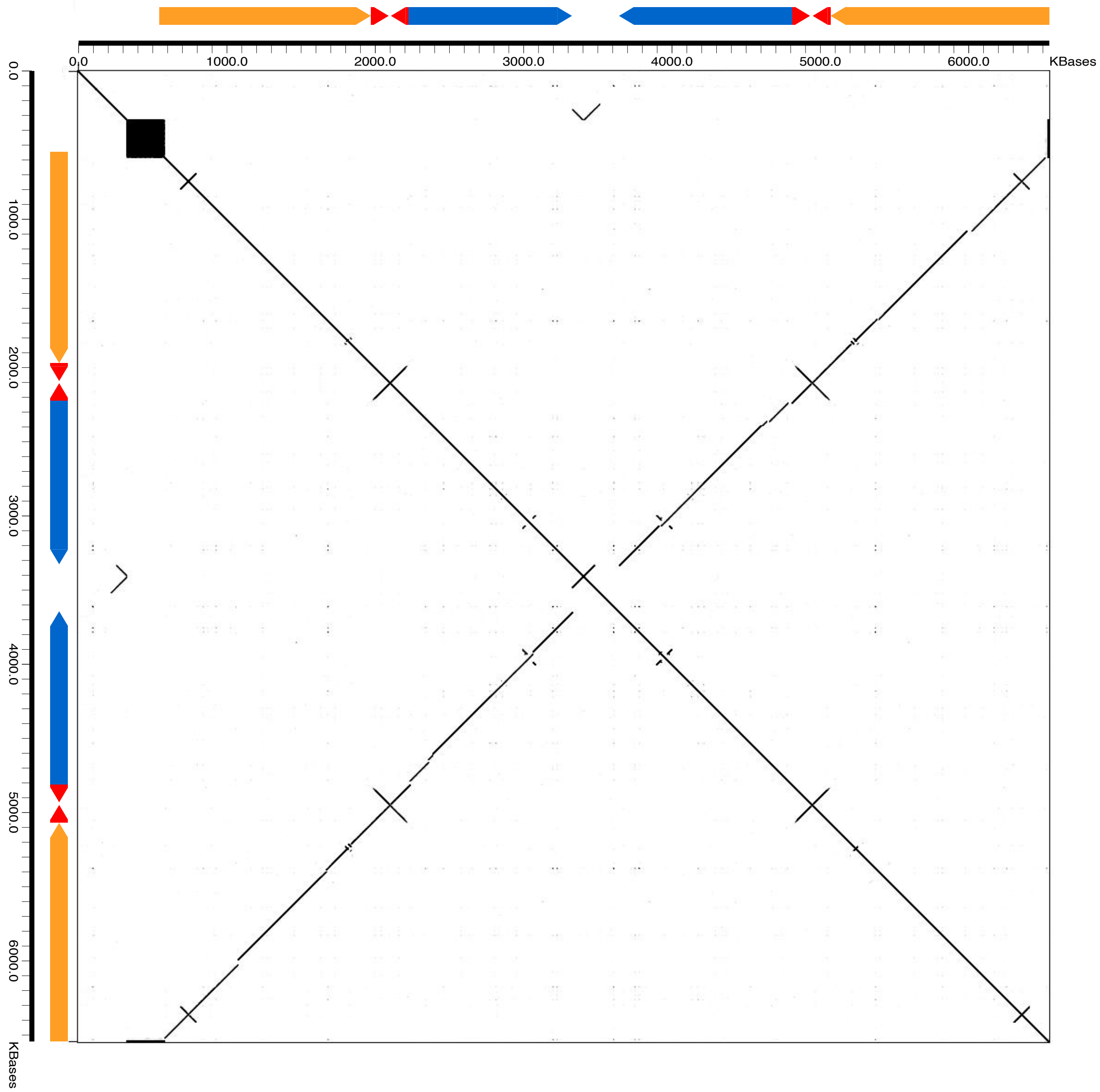


**Fig S2. Ampliconic structure of chrY (HG994382.1)**

A self-similarity dotplot of the mCanLor1.2 Y chromosome sequence drawn with a window size of 100 bp is shown. The colored arrows correspond to ampliconic sequences identified in the assembly. The entire sequence represents a complex palindromic structure.

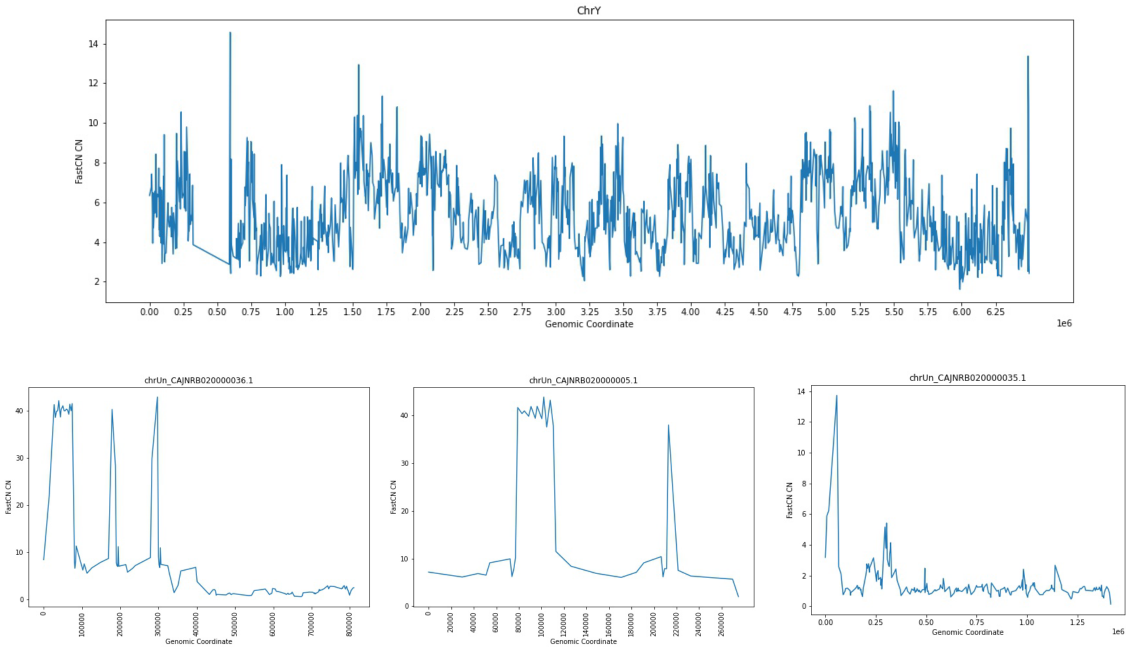


**Fig S3. Copy-number profile across Y-chromosome sequences**

Copy-number was inferred using fastCN applied to Illumina genome sequence data generated from the sample used to create the mCanLor1.2 assembly. Results are shown for the mCanLor1.2 Y-chromosome sequence (top) as well as three additional unplaced assembled contigs that are associated with the Y chromosome.

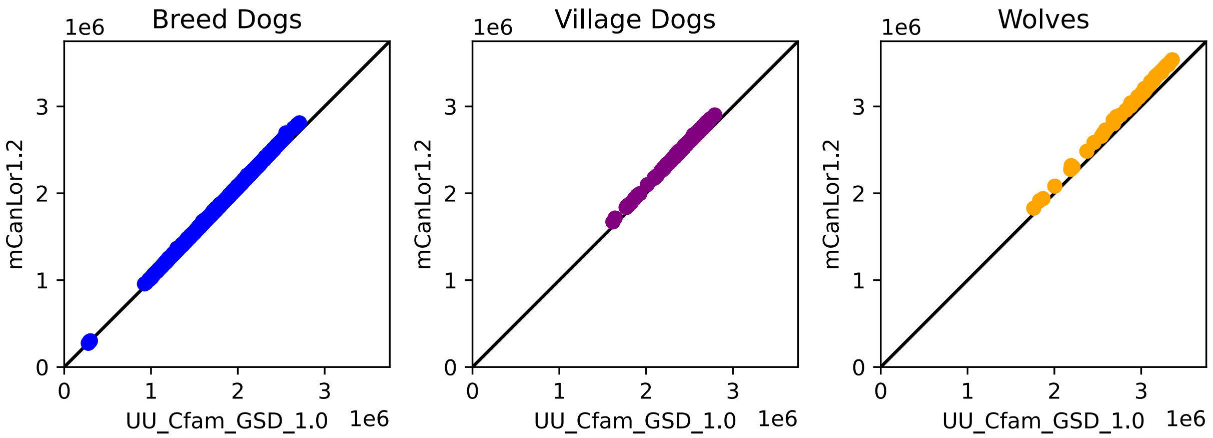


**Fig S4. Number of heterozygous sites identified based on alignment to UU_Cfam_GSD_1.0 and mCanLor1.2**

A scatter plot of the number of heterozygous sites identified from alignment to each reference genome is shown for breed dogs, village dogs, and wolves.

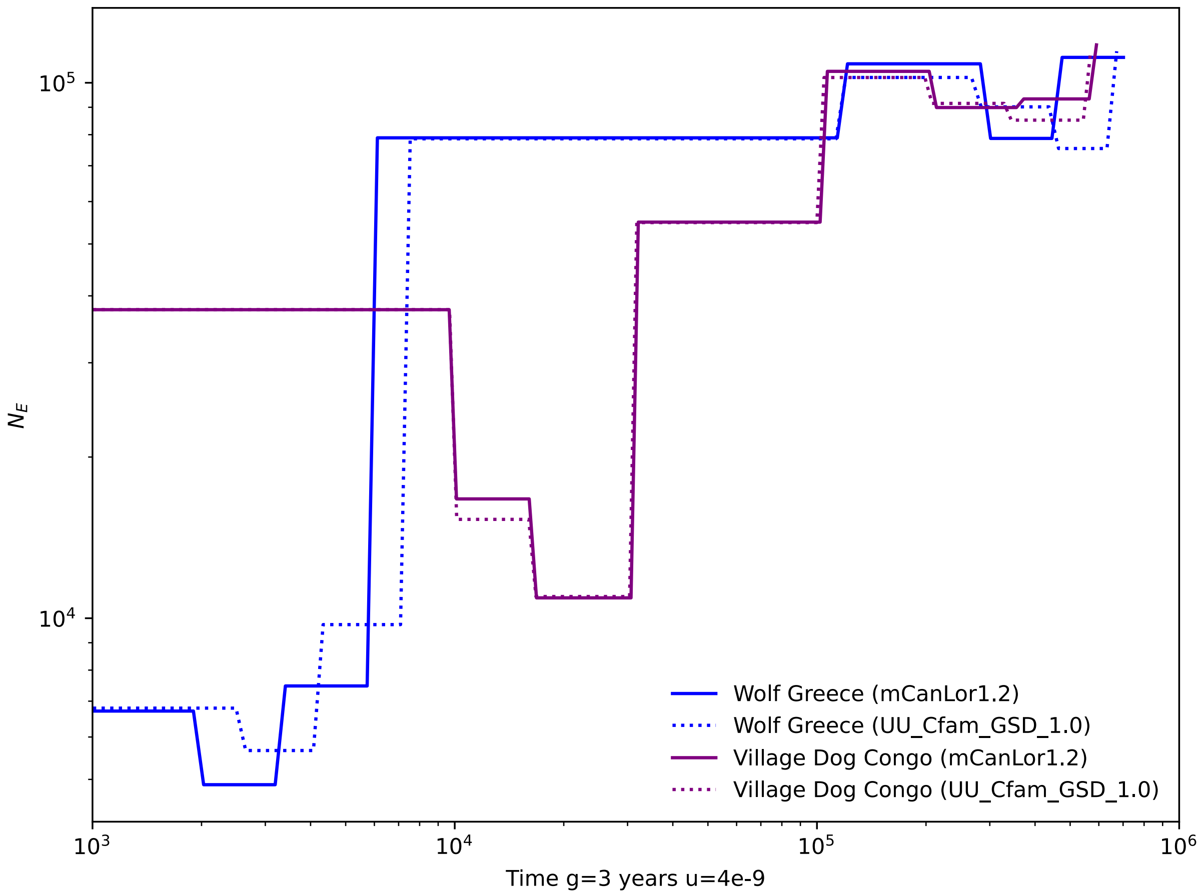


**Fig S5. Comparison of population size histories inferred using different reference assemblies.** The population history for wolves from Greece (blue lines) and village dogs from Congo (purple lines) was estimated using sequence alignment to the mCanLor1.2 (solid lines) and UU_Cfam_GSD_1.0 (dashed lines) assemblies. Inference was performed using SMC++.

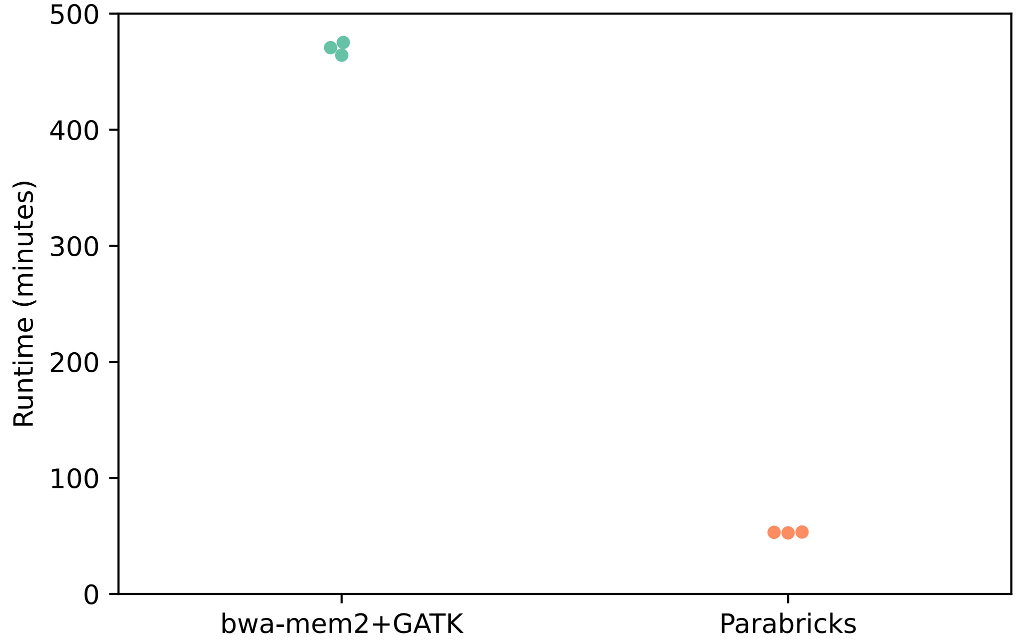


**Fig S6. Runtime comparison of standard and GPU analysis pipelines.**
Illumina fastq reads from sample ACKR000001 (SRR12330404) were processed against the UU_Cfam_GSD_1.0 assembly using a standard pipeline (left) and the GPU-enabled Parabricks pipeline (right). Total runtime (wall time) results are plotted from three analysis runs. Analysis includes alignment to the reference genome, sorting and duplication marking, base recalibration, and generation of GVCF-formatted variant files. The standard CPU pipeline ran on a single compute node with an attached solid-state drive (SSD) and used 24 3.0 GHz Intel Xeon Gold 6154 CPUs. The Parabricks pipeline ran on a single compute node and used two NVIDIA Tesla V100 GPUs and 24 2.4 GHz Intel Xeon Gold 6148 CPUs.

**Supplementary Table 1 Sequence deleted in UU_Cfam_GSD_1.0 that overlaps with annotated exons** (see excel file)
